# Astrocytic HIV-1 Nef expression decreases glutamate transporter expression in the nucleus accumbens and increases cocaine-seeking behavior in rats

**DOI:** 10.1101/2024.10.10.617598

**Authors:** Jessalyn Pla-Tenorio, Bethzaly Velazquez-Perez, Yainira Mendez-Borrero, Myrella Cruz-Rentas, Marian Sepulveda-Orengo, Richard J. Noel

## Abstract

Cocaine use disorder is an intersecting issue in populations with HIV-1, further exacerbating the clinical course of the disease, contributing to neurotoxicity and neuroinflammation. Cocaine and HIV neurotoxins play roles in neuronal damage during neuroHIV progression by disrupting glutamate homeostasis in the brain. Even with cART, HIV-1 Nef, an early viral protein expressed in approximately 1% of infected astrocytes, remains a key neurotoxin. This study investigates the relationship that exists between Nef, glutamate homeostasis, and cocaine in the NAc, a critical brain region associated with drug motivation and reward. Using a rat model, we compared the effects of astrocytic Nef and cocaine by molecular analysis of glutamate transporters in the NAc. We further conducted behavioral assessments for cocaine self-administration to evaluate cocaine-seeking behavior. Our findings indicate that both cocaine and Nef independently decrease the expression of the glutamate transporter GLT-1 in the NAc. Additionally, rats with astrocytic Nef expression exhibited increased cocaine-seeking behavior but demonstrated sex dependent molecular differences after behavioral paradigm. In conclusion, our results suggest the expression of Nef intensifies cocaine-induced alterations in glutamate homeostasis in the NAc, potentially underlying increased cocaine-seeking. Understanding these interactions better may inform therapeutic strategies for managing cocaine use disorder in HIV-infected individuals.

## 1. Introduction

HIV-1 remains a global epidemic that affects millions worldwide. In the United States alone, approximately 35,000 contract HIV yearly [1]. Prior to antiretroviral therapy, HIV infection caused an early death due to progression to AIDS; current therapeutic advances allow HIV-infected individuals a life-expectancy approaching that of non-infected [2]. Despite this great progress in the clinical management of HIV-1, new complications arise due to the chronicity of the infection [3]. In particular, it has increased the prevalence of HIV-associated neurocognitive disorders (HAND), a variety of conditions impairing cognitive function that arise due to the effects of the neurotropic virus in the central nervous system [4, 5]. To effectively address these challenges, it is important to study their interaction with other common public health issues in this affected community and how they can overlap to further alter neural function.

Substance use disorder is a major public health issue that intersects with HIV-1 infection and clinical management. Drug use has had a significant role in the global spread of HIV [6]. Cocaine use prevalence is higher in individuals who contract the virus, estimated to be ranging from 5% to 15% of HIV individuals as stated across multiple studies [7–9], which could be partially explained by the promotion of high-risk behavior due to drug effects [10, 11]. Not only does cocaine use disorder increase chances of acquiring the virus, it also hastens the progression of the disease by altering the body’s immune response as well as affecting the adherence to medication [12]. Thus, the comorbid use of cocaine in HIV-infected individuals can contribute to the vulnerability to and development of HAND; and contrarily can make it difficult to treat and overcome the substance use disorder itself [12, 13].

In this study, we explore the complex network of interactions between cocaine use disorder, HIV-1 and glutamate homeostasis in the nucleus accumbens (NAc), an area of the brain central to motivation, reward, and addiction. Neuronal homeostasis and function depend on astrocytes, the most prevalent type of glial cell in the CNS [14, 15]. Astrocytes are essential for preserving the microenvironment, controlling ion and neurotransmitter levels, and maintaining the blood brain barrier [15]. Astrocytes play a crucial role in regulating glutamate levels in the brain by up-taking excess glutamate from the synaptic cleft, preventing excitotoxicity, converting glutamate into non-toxic substances or recycling it to presynaptic terminals [14]. Glutamate itself plays an important role in synaptic transmission, memory, learning, and neural communication. In the NAc particularly, glutamate plays a key role in mediating reward and pleasure, influencing motivation, and contributing to addictive behaviors [16].

Astrocytes are susceptible to HIV-1 infection, and although they do not produce viral progeny, infected astrocytes produce early viral proteins of which the most abundant is Nef [17, 18]. Notably, antiretroviral therapy does not target HIV-1 transcription and thus does not prevent early protein synthesis [19]. The intracellular expression of Nef has been found to interact with signaling pathways leading to oxidative stress, triggering inflammation, increasing neural degeneration, and dysregulating astrocyte function [17, 20–23]. Moreover, the presence of Nef has shown to decrease glutamate clearance, leading to excitotoxicity [17, 24]. These myriad interactions disrupt homeostasis and contribute to the overall neurotoxicity of the virus [17].

On the other hand, cocaine has shown to increase expression of GFAP in astrocytes, suggesting cocaine also takes a role in promoting astrogliosis and disturbing the brain homeostasis [25]. In the NAc, cocaine-use results in synaptic potentiation and stimulates glutamate release brought on by conditioned signals and drug exposure [25, 26]. Through this excess of glutamate and access to glutamate postsynaptic receptors, this potentiated release promotes the synaptic plasticity that underlies cocaine-seeking behavior [16, 25, 26].

Glutamate homeostasis is tightly regulated by a family of transporters, and alterations to this equilibrium have been linked to several neurological and psychiatric diseases, including addiction [14]. We seek to understand the intricate interplay through which HIV-1 Nef and cocaine can disturb this homeostasis and contribute to the higher prevalence of cocaine addiction in individuals living with HIV. We hypothesized that HIV-1 Nef expressed in astrocytes of the NAc would cause higher cocaine seeking behavior in rats through changes in glutamate regulation and stronger synaptic transmission in the reward pathway.. A better understanding of the intersection between cocaine and HIV-1 neuropathogenesis can lead to possible changes and advancements in the therapeutic approach to HIV-infected cocaine users.

## 2. Materials and Methods

### 2.1 Animals

Adult 60 to 70-day male and female Sprague Dawley rats were obtained from the Ponce Health Sciences University colony. Animals were separated and single housed. Animals were handled for 5 days prior to beginning any procedure and were maintained under standard conditions with a 12h/ light/dark cycle with free access to food and water for 2.3. Animals for 2.5 and 2.6 were kept at an inverted 12h/dark/light cycle with an 18g food diet. All animal work was approved by the Ponce Health Sciences University Institutional Animal Care and Use Committee (IACUC) and followed the National Institutes of Health’s Guide for the Care and Use of Laboratory Animals.

### 2.2 Bilateral Stereotaxic Surgery

At the time of surgery, adult male rats (∼275-310g) and female (∼200-235g) animals were placed on a stereotaxic apparatus and anesthetized with isofluorane (3-4%). Then they were stereotaxically microinjected in the targeted area of the nucleus accumbens core, NAc core. Bilateral coordinates (mm) for males: +1.5. anterior/posterior, +2.6 medial/lateral, -7.4 dorsal/ventral and +1.5. anterior/posterior, -2.6 medial/lateral, -7.4 dorsal/ventral. Bilateral coordinates (mm) for females: +1.5. anterior/posterior, +2.6 medial/lateral, -7.2 dorsal/ventral and +1.5. anterior/posterior, -2.6 medial/lateral, -7.2 dorsal/ventral. Infusion was the packaged lentivirus vector of Lenti-GFAP-Nef-IRES-mCherry (Vector Builder cat #LVMP(VB230201-1721wex)-K2) to express HIV-1 Nef and an enhanced mCherry fluorescent protein driven by the astrocyte specific GFAP promoter, or Lenti-GFAP-IRES-mCherry for controls. Viral vector stocks were independently aliquoted and stored at -80°C until use and thawed in ice. Virus was microinjected bilaterally to the NAc core using 26-gauge cannulas (Plastics One, Roanoke, VA) in a single and dual configuration (1 ul per injection site, 0.05 ul/min injection rate) and left to diffuse for 10 minutes. After diffusion, the microinjector was slowly removed over 1–2 min as described previously by Testen et al., 2018 [27]. Then, the incision site was sealed with stitches and rats were injected with analgesic ketorolac (10mg/kg i.p.). Animals were provided seven days to recover from surgery and five weeks after infusion to begin experimental procedures. After molecular and behavioral experiments, all infusion sites were histologically confirmed. Animals with improper infusion placements or significant diffusion outside of the target region were rejected.

### 2.3 Cocaine injection procedure

Cocaine hydrochloride (provided by the NIDA drug supply program) was dissolved in sterile saline (0.9% solution) to a concentration of 15mg/mL. For drug administration, animals were injected i.p. at the same time with saline or cocaine. This specific dose is consistent with other literature reviews [28–32]. After 24 hours, the animals were anesthetized with an injection of pentobarbital (65 mg/kg i.p.). When the animal was deeply anesthetized (without response to tail pinch), the thoracic cavity was cut with scissors to expose the heart. The right atrium was cut, a dull needle was inserted into the left ventricle, and 20-30 mL of saline solution was perfused to remove the blood from the brain. Immediately following, the animal was decapitated with guillotine and the brain was divided in half. The brains were cryopreserved for further molecular analyses of the NAc. Vaginal smears of female rats were taken at sacrifice for corroboration of estrus cycle phase.

### 2.4 Immunofluorescence

For analysis of GFAP, GLT-1 and xCT expression by immunofluorescence staining, 30um brain sections were cut in a cryostat (LeicaCM1520) and placed in a 24 well plate, one section per well. Then, sections were washed three times in PBS for 10 minutes each, followed by two hours incubation in 10% normal goat serum, 0.3% Triton X-100 in PBS. The tissues were then incubated overnight in primary antibodies GFAP (dil. 1/200, cat#644701, Biolegend), EAAT2/GLT-1 (dil. 1/400, cat#AB1783, Millipore) and SCL711A/xCT (1/200 dil., cat# PA1-16893, Invitrogen), respectively. A tissue section was used as a negative control receiving PBS instead of primary antibody. The second day, the tissues were washed three times for 10 minutes in PBS, followed by one hour incubation in Alexa fluor 488 goat anti-mouse secondary antibody (1/400 dil., cat# A11029, Invitrogen), Alexa flour goat anti-guinea pig secondary antibody (1/400 dil., cat#, supplier) or Alexa fluor 594 goat anti-rabbit secondary antibody (1/200 dil., cat# A11037, Invitrogen), respectively. After washing three times with PBS for 10 minutes each, sections were incubated for 5 minutes in DAPI (R37606: Invitrogen). Tissues were washed in three intervals of 5 minutes with PBS buffer and then mounted on slides using ProLong Gold Antifade mountant (P36934, Invitrogen). Three representative areas were photographed for each brain section using a Nikon Confocal microscope at 60X magnification and the images were further analyzed using ImageJ software.

### 2.5 Catheter Implantation Surgery

All rats were given a 5-week recovery period from infusion surgery. After recovery, rats were food deprived for a 24-hour period prior to receiving food training sessions in the operant chambers consisting in acquisition of sucrose pellets through the operant behavior of lever pressing. The operant chamber had two levers: an active lever that provided a sucrose pellet to the rat and an inactive lever that had no effect. Once 100 presses were reached, the rats finished the food training session. After training, animals were exposed to 1.5-2.5% of isoflurane plus Ketorolac (10mg/kg) as an analgesic for surgical procedures. For implantation of the catheter, a guide cannula (Plastics One, cat# C313G) was attached to a silastic tube (.025 ID, .047 OD Bio-sill), inserted subcutaneously between the shoulder blades, and exited through the skin via a dermal biopsy hole (3mm). The catheters were then secured by subdermal surgical mesh (Atrium) and the cannula exiting the skin. The other end of the catheter was inserted 3cm into the right jugular vein and then securely sutured to the underlying muscle tissue. A catheter cap was used when the rats were not connected to infusion pumps. Immediately following catheterization, animals were provided seven days to recover from surgery. After surgery, the IV catheters were flushed daily with 0.1 ml each of 5 mg/ml gentamicin and 70 U/ml heparinized saline for five days following surgery to maintain catheter patency.

### 2.6 Self-Administration Procedure

Patency of the intravenous catheter was maintained by daily infusions of 0.1mL of heparin saline (70 units/ml) and checked periodically by infusion of 1mg/0.1mL, i.v. of Propofol (1%; Abbot), a fast-acting anesthetic delivered through the catheter that causes immediate loss of muscle tone and brief sedation (∼5 min). Animals were excluded from the experiment if they failed the propofol test. Rats were placed in a self-administration apparatus and allowed to lever press for cocaine on a FR1 schedule of reinforcement (two hours per day for 10 days). Each self-administration cage contained one active and one inactive lever. All sessions would begin and end between the hours of 0700 and 1100 (light-cycle). Cocaine (5 mg/mL in 0.9% sterile saline) was housed in a syringe pump external to the training chamber. Each press of the active lever delivered approximately 0.1mg/kg (in 0.020 ml total volume/infusion for 300 g rat) [33]. A cue light over the active lever was illuminated and a tone (200Hz/70db/5 seconds) was delivered at the initiation of each infusion. For all experiments, the active lever was deactivated for a 20 second period after each infusion, although all responses on both the active and inactive levers were recorded. Extinction phase consisted of rats being placed inside the apparatus without cocaine infusions, cue light or tone, for 14 consecutive days. A cue-primed reinstatement test was given the day after the last extinction, where rats were re-exposed to the cocaine associated cues (light and tone) with a right (active) lever press. Two more days of extinction sessions were given between the two tests. For cocaine-primed reinstatement phase rats received one dose of 10mg/kg i.p. cocaine or saline immediately before placing them in respective operant chambers without cues or infusions. Rats were sacrificed 24 hours after cocaine-primed reinstatement as described in a previous section. Vaginal smears of female rats were taken at the beginning of each behavioral phase (SA Day 1 and Ext Day 1), before each test (Cue and Cocaine-Primed Reinstatement) and at the day of sacrifice for corroboration of estrus cycle phase.

### 2.7 Western blotting

NAc tissue punches were lysed using RIPA buffer of 150mM sodium chloride, 1.0% NP-40, 0.5% sodium deoxycholate, 0.1% sodium dodecyl sulphate and 50mM Tris, pH 8.0. Protein concentration was determined using Micro BCA Protein Assay Kit (Thermo Scientific, cat# 23235). An equal amount of protein (25ug) was assessed by SDS-PAGE in ANY KD gel (BioRad, cat# 4569033) and transferred into a polyvinylidene difluoride membrane (PVDF) (Bio-Rad, cat# 162-0175). Membranes were blocked with 5% Bovine serum albumin (Sigma, cat# B4287) in Tris-Buffered Saline Tween at room temperature for 4 hours and incubated overnight at 4°C with primary antibodies. The following antibodies were used: EAAT2/GLT-1 1:5000 (cat#AB1783, Millipore), SCL711A/xCT 1:1000 (Invitrogen, cat# PA1-16893), Anti-HIV-Nef monoclonal 1/250. (NIH AIDS Reagent program, cat# 3689), β-actin 1:5000 (Sigma, cat# A5441), GFAP 1:1000 (cat#644701, Biolegend). Then, gels were washed 2 times for 10 minutes each with TBST, and 2 times for 10 minutes with Blocking solution (non-fat dry milk) and incubated with secondary antibodies Rabbit 1:5000 (ECL anti-Rabbit IgG, peroxidase-linked species-specific whole antibody, cat# NA934V) or Mouse 1:5000 (ECL anti-Mouse IgG, perox-idase-linked species-specific whole antibody, cat# NA931V) for 1 hour at room temperature. Blots were detected using Super Signal Chemiluminescence Substrate (Thermo Scientific, cat# 34578) using a ChemiDoc XRS+ Imaging system (Bio-Rad). All western blots were from separate experiments, each including all four treatment groups to reduce the impact of experimental variability. Quantification of the western blots were done using Image J 1.54d (National Institute of Health, USA). Band densities were normalized to β-actin from the same sample and blot to control for loading. All western blot data are presented as protein expression compared to control.

### 2.8 Nef Free-floating Immunohistochemistry

Brain sections were cut at 30µm thickness with a cryostat (Leica CM 1520) and placed in a 24 well plate, one section per well. This was followed by a 3% Hydrogen peroxide (Sigma-Aldrich) incubation for 15 minutes to block endogenous peroxidase and three PBS 1X 3% Triton washes, 15 minutes per wash. Sections were blocked with normal goat serum (BioGenex, cat#HK112-9KE) for 1 hour followed by an overnight incubation with Nef primary antibody (NIH AIDS Reagent program, cat# 3689; 1/50 dil.). A negative control with PBS instead of primary antibody was run in the procedure. On the second day, the sections were rinsed with three changes of PBS 1X 0.3% Triton for 15 minutes each, followed by a 4°C overnight MultiLink secondary antibody incubation (Super Sensitive Link-Label IHC Detection System, cat#LP000-UL, 1/20 dil. in PBS, BioGenex, Fremont, CA, USA). On the third day, the sections were rinsed with three changes of PBS 1X for 15 minutes each, followed by 2 hours Streptavidin Peroxidase (Super Sensitive Link-Label IHC Detection System, cat#LA000-ULE, 1/20 dil. in PBS, BioGenex, Fremont, CA, USA). The sections were rinsed with three washes of PBS 1X for 15 minutes each, followed by 3,3’-Diaminobencidine (DAB) incubation (BioGenex, cat# HK542-XAKE, Fremont, CA, USA) on each section and the exposure was monitored for 30 seconds. Then, the sections were counterstained with hematoxylin and mounted on slides for confocal microscope visualization of Nef expression. Two representative areas were photographed at high power field for each tissue for further analysis. (Nikon Confocal Microscope, 60X magnification).

### 2.9 Data analysis

Statistical analysis was performed in GraphPad Prism using a One-way ANOVA with Tukey Post-Hoc with multiple comparisons analysis to compare groups (Figures 2, 3, and 7). Male and female groups molecular analysis of figures 2 and 3 were combined since no changes were observed between sex. For behavioral analysis (figures 5 and 6), Two-way ANOVA with uncorrected Fischer’s LSD Post-Hoc analysis was done to compare groups per day. Values were reported as the mean ± the standard error of the mean (SEM).

**Figure 1.**
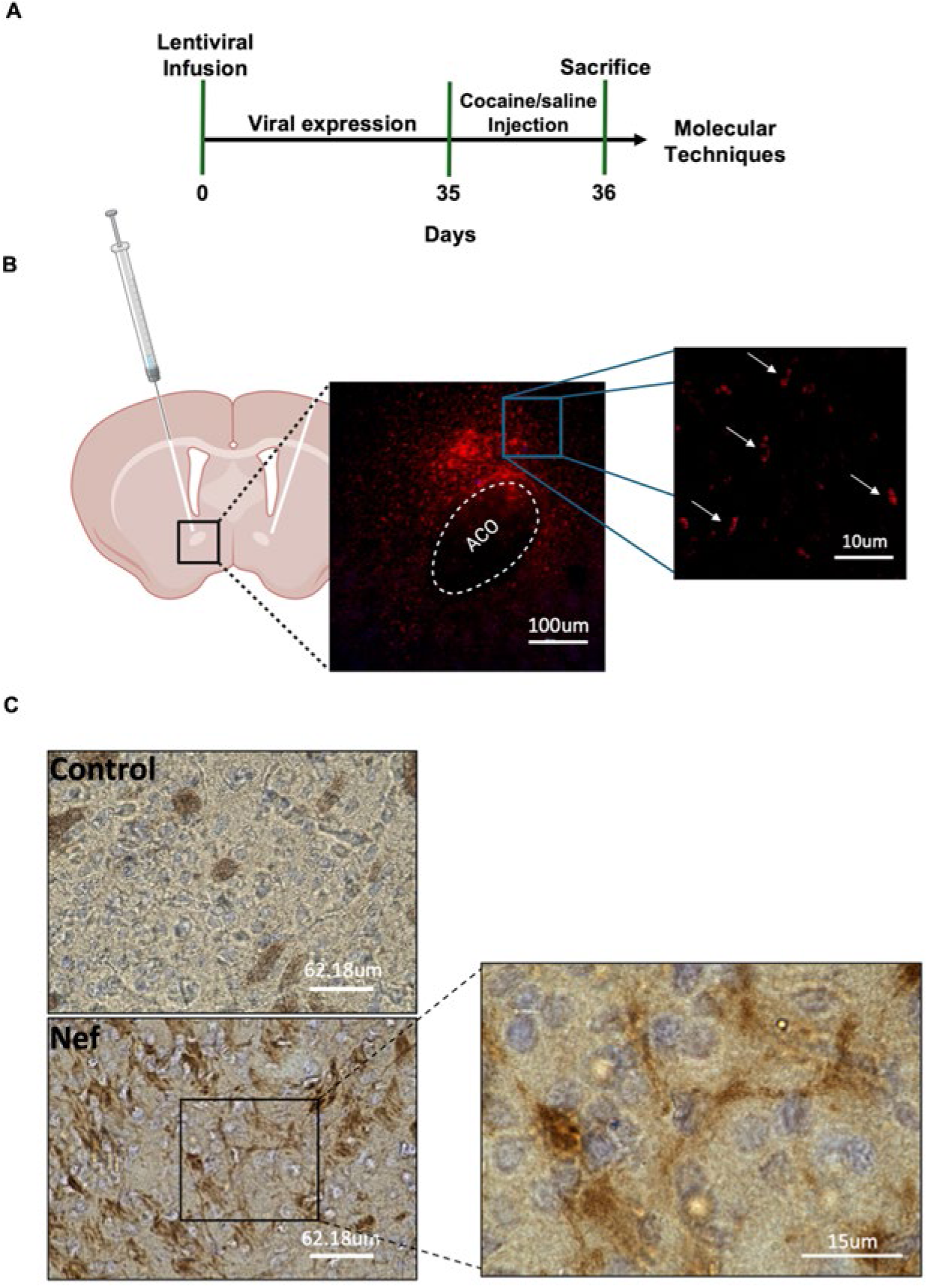
Timeline and schematic representation of bilateral infusions in the NAc at 5 weeks. (**A**) Specific timeline used for molecular experiments with rats. (**B**) Schematic representation showing location of the bilateral infusion site on the corresponding coronal section (red dots). Viral expression spans from approximately bregma 0.70mm to 1.60mm. GFAP specific promoter expression of mCherry in NAc of Sprague Dawley rats 5 weeks after infusion. 10x magnification (100um scale). Right image at 100x magnification (10um scale) shows positive punctate cytoplasmic staining in the NAc (right hemisphere). (**C**) Representative images of immunohistochemical staining with Nef antibody (1:50) in NAc slice of Control (top) and Nef treated (bottom) rat brain tissue. DAB staining in brown with hematoxylin for nuclear counterstain. Pictures of 30um thick tissues taken at 60x magnification (62.18um scale). Zoomed area at 100x magnification (15um scale) of Nef positive NAc slice demonstrating DAB staining with an astrocyte morphology.

**Figure 2.**
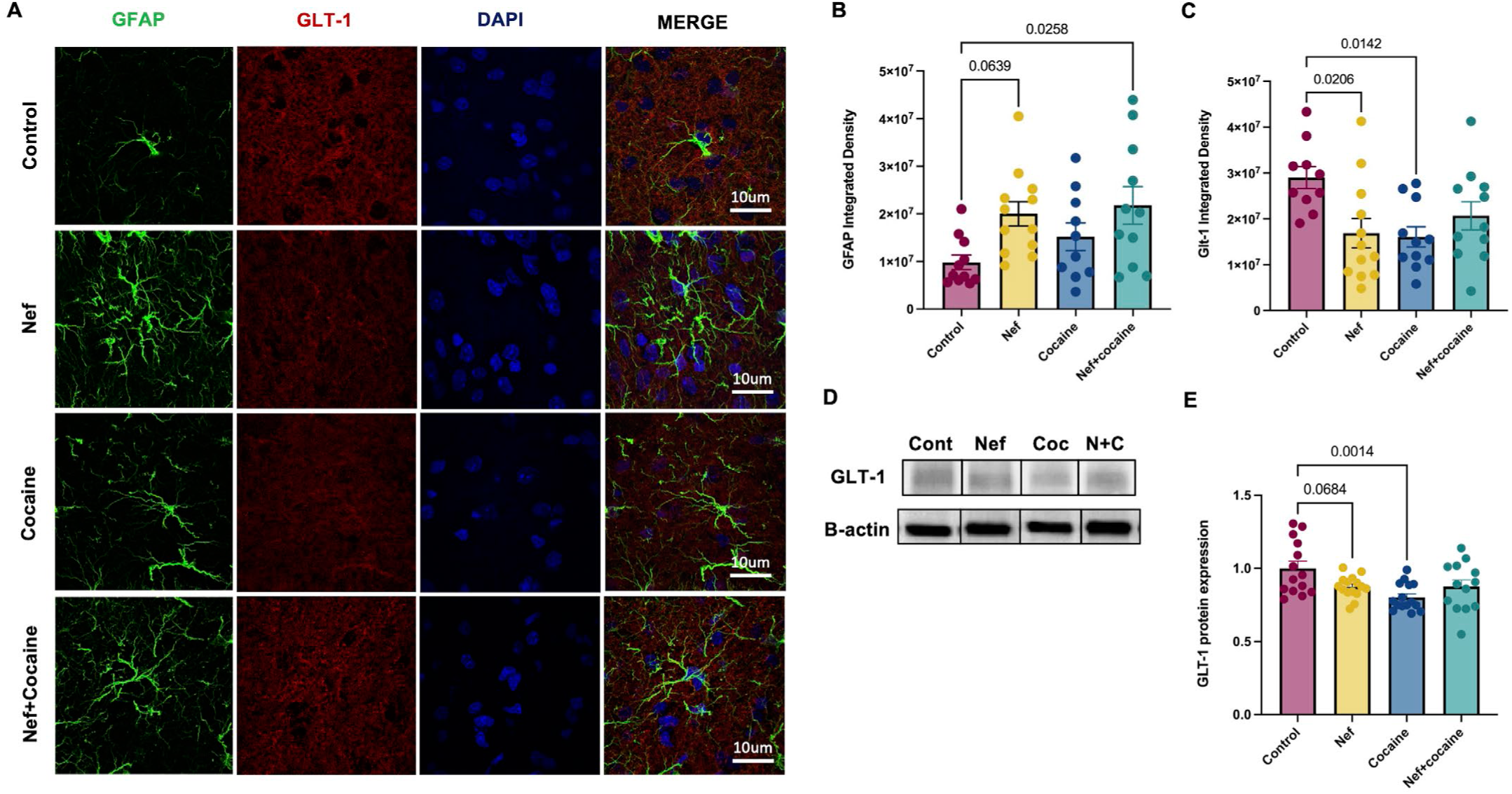
Nef expression in the NAc combined with cocaine causes an increase in GFAP. (**A**) Representative images of free-floating immunofluorescence to identify astrocyte expression using GFAP (green, 1:200) and GLT-1 (red, 1:400). Nuclear counterstaining was done using DAPI. Pictures were taken at 100x magnification (10um scale). (**B**) Integrated density of GFAP in the infusion site demonstrates increased expression in the Nef+cocaine treatment. (**C**) Integrated density of GLT-1 in the infusion site shows decreased expression after 24 hours of cocaine injection (15mg/kg, i.p.) in Nef and cocaine treatments compared to control. Each sample represents an average of 3-4 60x mag fields per rat in each treatment. (**D**) Representative protein bands of GLT-1 (65kD) and 𝛽𝛽-actin (42kD). From left to right: Control, Nef, Cocaine, Nef+cocaine. (**E**) Protein expression of GLT-1 shows decrease 24 hours after cocaine injection (i.p.). Ordinary one-way ANOVA with Tukey Post-hoc and multiple comparisons was used to compare data within groups (Control: n=10-14; Nef: n=12-15; Cocaine: n=10-15; Nef+cocaine: n=11-13). Data represent mean +/- SEM adjusted to 𝛽𝛽-actin and normalized to control.

**Figure 3.**
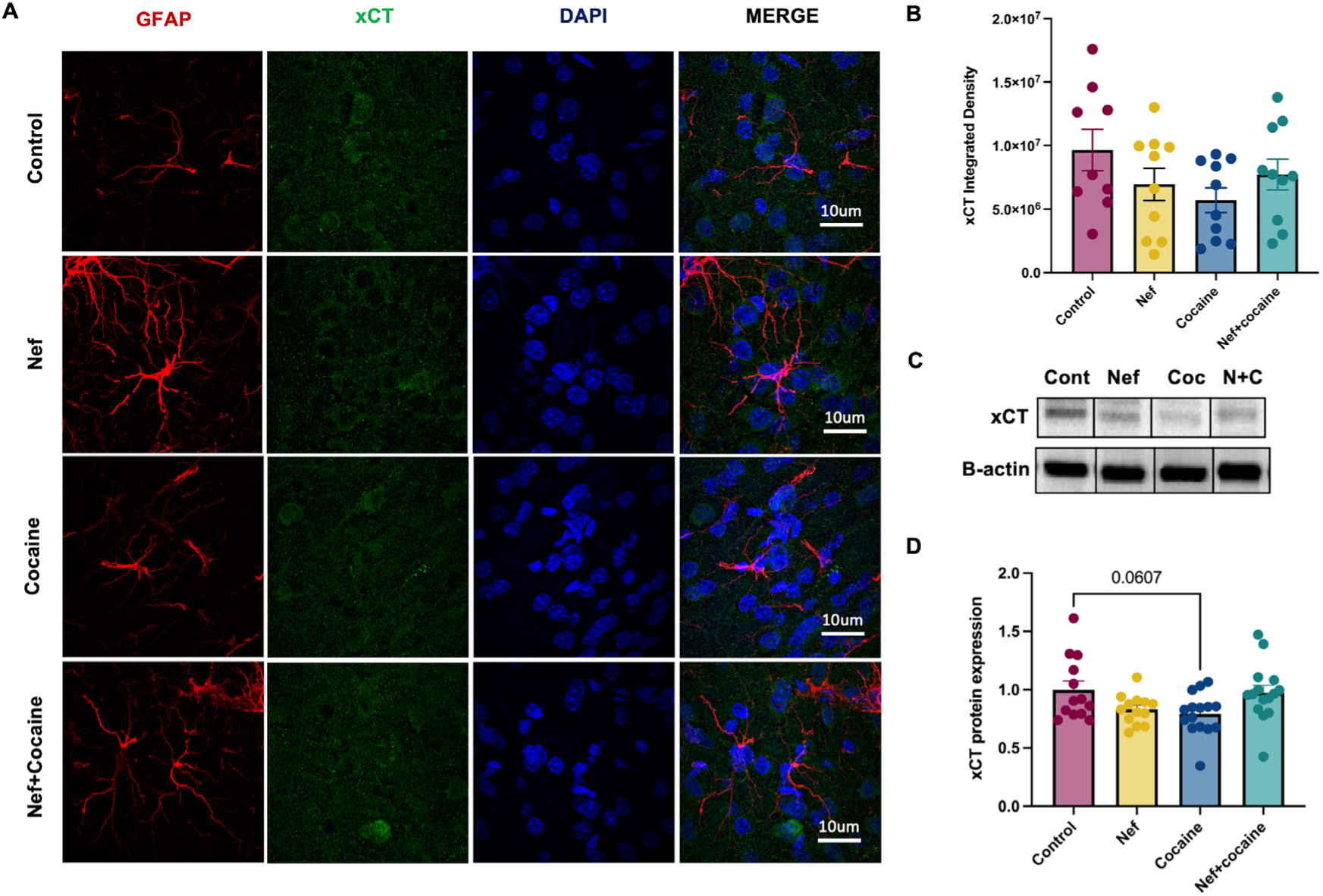
Nef expression in the NAc causes no effect on xCT expression. (**A**) Representative images of free-floating immunofluorescence to identify astrocyte expression using GFAP (red, 1:200) and xCT (green, 1:200). Nuclear counterstaining was done using DAPI. Pictures were taken at 100x (10um scale). Each sample represents an average of 3-4 60x mag fields per rat in each treatment. (**B**) Integrated density of xCT expression in the NAc shows no significant differences between treatments. (**C**) Representative protein bands of xCT (37kD) and 𝛽𝛽-actin (42kD) per treatment. From left to right: Control, Nef, Cocaine, Nef+cocaine. (**D**) Protein expression of xCT in nucleus accumbens shows decreased trend in cocaine treatment compared to control. Ordinary one-way ANOVA with Tukey Post-hoc and multiple comparisons was used to compare data within groups (Control: n=9-13; Nef: n=10-13; Cocaine: n=10-15; Nef+cocaine: n=10-15). Data represent mean +/- SEM adjusted to 𝛽𝛽-actin and normalized to control.

**Figure 4.**
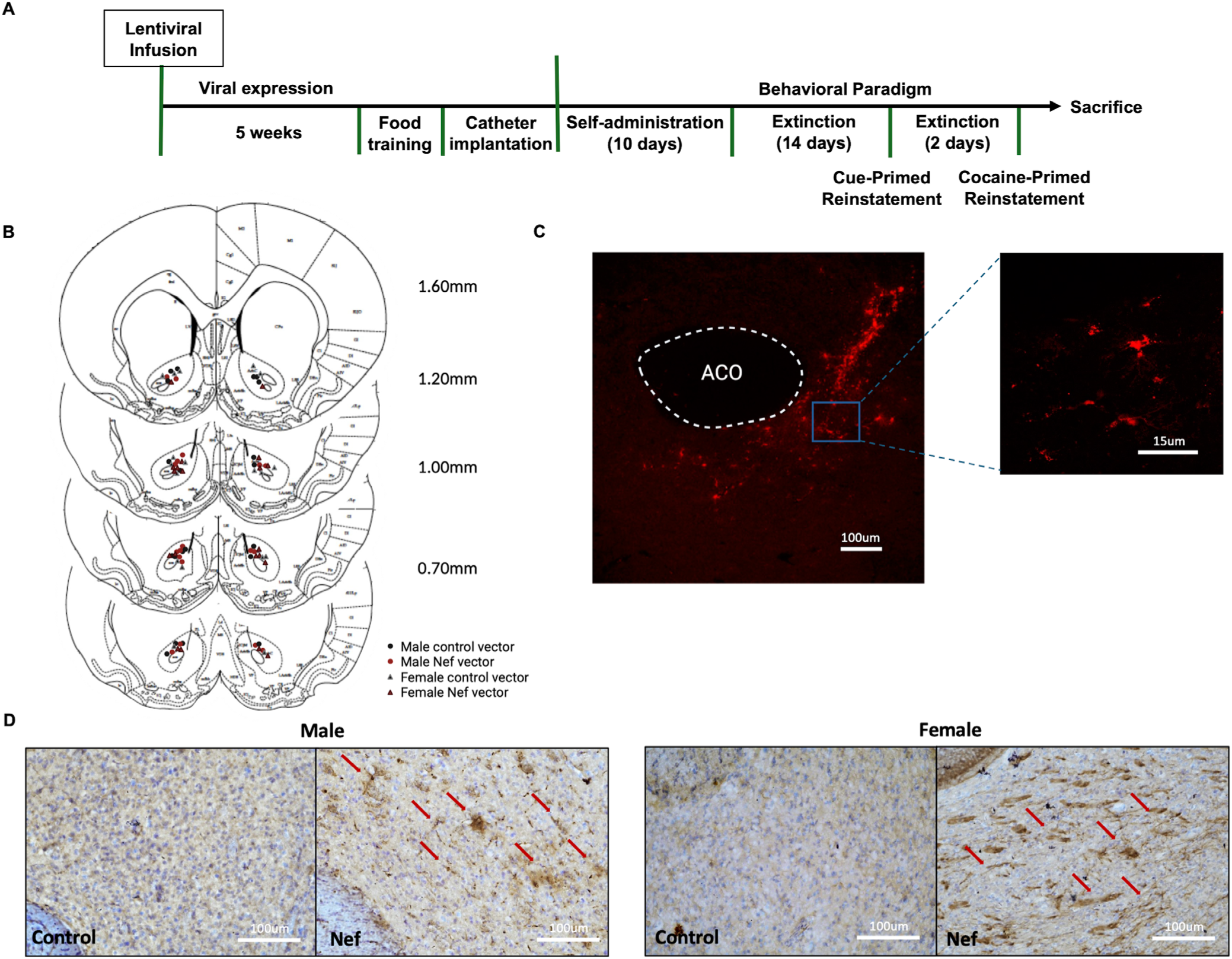
Behavioral timeline and schematic representation of lentiviral infusions in NAc core. (**A**) Behavioral timeline used for cocaine self-administration experiments with rats. (**B**) Schematic representation showing placement of the infusion site of each rat in the corresponding coronal section of the NAc ranging from 0.70mm to 1.60mm (Male control vectors - black dots; Male Nef vectors - red dots; Female control vectors - red triangles; Female Nef vectors - gray triangles). (**C**) GFAP specific promoter expression of mCherry in NAc of Sprague Dawley rats 10 weeks after infusion. 10x magnification (100um scale) with zoomed in area at 15um scale showing morphology of cells infected, respectively. (**D**) Representative images of Nef immunohistochemistry at infusion site after behavioral paradigm. DAB staining in brown with hematoxylin for nuclear counterstain. Red arrows represent astrocytes positive for Nef. Pictures of 30um thick tissues taken at 20x magnification (100um scale).

**Figure 5.**
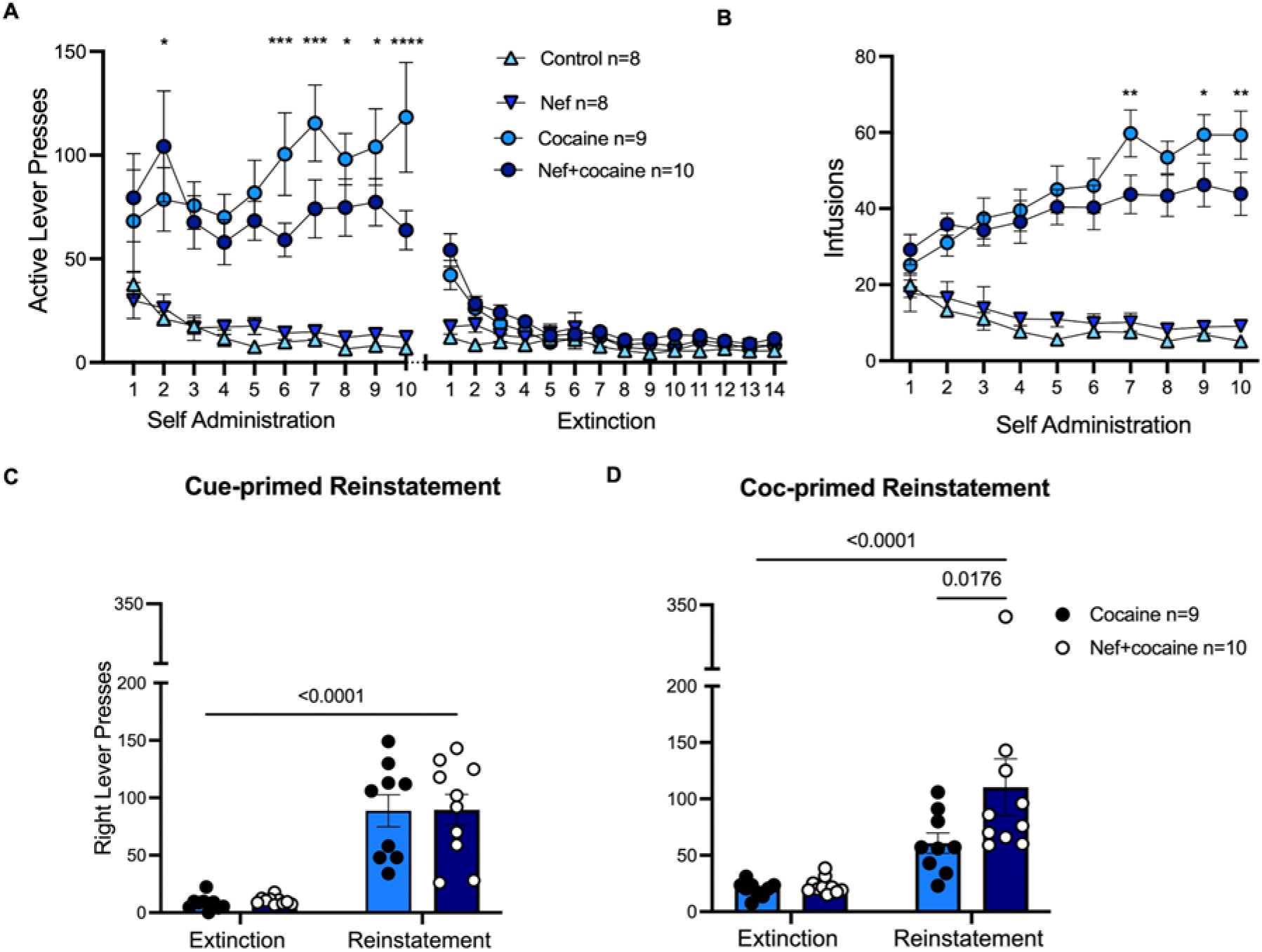
Nef-expressing male rats show more cocaine seeking behavior than Cocaine male rats. (**A**) Active lever presses of Control, Nef, Cocaine, and Nef+cocaine male rats during self-administration phase and extinction. (**B**) Cocaine infusions of Cocaine vs Nef+cocaine rats and saline infusions for Control and Nef rats during self-administration. *, **, ***, **** represent significant differences between Cocaine and Nef+cocaine groups during self-administration. P < 0.05, 0.01, 0.001, 0.0001, respectively. (**C**) Average right lever presses (correct) during extinction of Cocaine vs Nef+cocaine rats compared to right active lever presses during cue-primed reinstatement. (**D**) Average right lever presses of Cocaine vs Nef+cocaine animals compared to right active lever presses during cocaine-primed reinstatement. Nef+cocaine rats significantly increased active lever presses in cocaine primed reinstatement compared to Cocaine group. Two-way, repeated measures ANOVA with Fisher’s LSD Post-Hoc analysis indicated all rats reinstated cocaine seeking compared with extinction responding. Data represent mean +/- SEM.

## 3. Results

### 3.1 Astrocytic Nef and cocaine had independent effects on glutamate transporter expression in the NAc

In this study, we proposed to test the hypothesis that with Nef present, one injection of cocaine is enough to produce strong neurophysiological changes in the NAc and alter the glutamate cycle. To address this question, we infused a lentiviral vector expressing Nef or mCherry into Sprague Dawley rats’ brains, specifically into the NAc core by bilateral stereotaxic surgery (Figure 1A). After 5 weeks, we verified the infusion site and expression of the vector in all rats after sacrifice. The control viral vector was found to positively express mCherry only in astro- cytes using the cell-specific GFAP promoter. Figure 1B shows a punctate mCherry expression pattern in astrocytes of the NAc core but not in areas outside of the infusion. To demonstrate Nef expression, we conducted an immunohistochemical staining using Nef antibody which showed positive DAB staining in astrocytes of the NAc core(Figure 1C).

We next tested whether Nef expression in astrocytes alone had direct effects on glutamate transporters and if a single cocaine exposure potentiated this effect. We first tested the expression of GLT-1, which is the most prevalent transporter to take up extracellular glutamate from the synaptic cleft [34]. Immunofluorescence analysis (Figure 2A) revealed that exposure to Nef in astrocytes decreases both the expression and intensity of GLT-1 fluorescence in the NAc core. We confirmed the expression of astrocytes in the area using the astrocyte marker, GFAP (Figure 2B) and demonstrated a clear downregulation of GLT-1 expression around reactive astrocytes (One-way ANOVA main effect between treatments: F(3,40)=4.231, P=0.0109; Tukey’s Post-Hoc with multiple comparisons: Control vs Nef: p=0.0206; Control vs. Cocaine: p=0.0142; Control vs. Nef+cocaine: p=0.1873; Figure 2C). However, while either cocaine or Nef alone decreased GLT-1 expression, the effect was not additive in combination. These immunofluorescence results were confirmed by western blot analysis, showing a trending decrease in GLT-1 transporter expression in NAc punches with Nef compared to brains without Nef or a single dose of cocaine (One-way ANOVA main effect between treatments: F(3,53)=5.286; P=0.0029; Tukey’s Post-Hoc with multiple comparisons: Control vs Nef: p=0.0684; Control vs Cocaine: p=0.0014; Control vs Nef+cocaine: p=0.0913; Figure 2D,E). Our results also demonstrate a significant increase in astrogliosis with Nef+cocaine compared to control. However, even if not significant, we also observed a trending increase in GFAP with the expression of Nef in the tissue compared to control (One-way ANOVA main effect between treatments: F(3,40)=3.548, P=0.0228; Tukey’s Post-Hoc with multiple comparisons: Control vs Nef: p=0.0639; Control vs Cocaine: p=0.5737; Control vs Nef+cocaine: p=0.0258; Figure 2B). This data demonstrates that astrocytic Nef expression has direct effects, not only on the reactivation of astrocytes in the brain, but also on their function. The effects of Nef expression in astrocytes in the NAc also seem to be similar to the physiological effects in the brain after a first exposure to cocaine.

We also measured expression of the cysteine-glutamate exchanger due to its importance in astrocyte maintenance of glutamate homeostasis. Past studies have demonstrated that this antiporter, also known as system Xc- or its light chain, xCT, is also downregulated in astrocytes in response to cocaine consumption [35, 36]. Here we sought to determine if this transporter is affected by Nef expression in astrocytes. Immunofluorescence analysis demonstrated no change in xCT integrated density after either exposure to cocaine or Nef in the NAc core (One-way ANOVA no main effect between treatments: F(3,35)=1.626, P=0.2009; Tukey’s Post-Hoc with multiple comparisons: Control vs Nef: p=0.4627; Control vs Cocaine: p=0.1552; Control vs Nef+cocaine: p=0.7200; Figure 3B). The effect, like with GLT-1, was not additive when exposed to both cocaine and Nef. Western blot analysis demonstrated only a trend towards a decrease after one dose of cocaine compared to control and no effect when exposed to Nef or Nef+cocaine (One-way ANOVA main effect between treatments: F(3,52)=3.293, P=0.0276; Tukey’s Post-Hoc multiple comparisons: Control vs Nef: p=0.1929; Control vs Cocaine: p=0.0607; Control vs Nef+cocaine: p=0.9886; Figure 3C, D). This data demonstrates that Nef expression in astrocytes in the NAc core does not affect xCT specifically measured at 5 weeks post infusion.

### 3.2 Male and female rats with Nef in the NAc responded differently to cocaine self-administration but resulted in higher cocaine-primed reinstatement

As mentioned before, cocaine use prevalence is higher in HIV-infected patients [7–9]. However, very few studies are focused on the long-term effects cocaine may have in these patients, even when the viral load is undetectable and HIV viral proteins, like Nef, are still present in the brain. In this section, we focus on how HIV-1 Nef can further disrupt the consequences (or physiological response) caused by an acute, long-term consumption of cocaine after a self-administration paradigm. To do this, we once again took Sprague Dawley rats, but this time exposed them to a cocaine self-administration paradigm after expression of the lentiviral vector (Timeline shown in Figure 4A). All rats were checked for lentiviral infusion placement and expression after sacrifice and if the vector was still expressed (Figure 4B, C). We also wanted to confirm that Nef was still present in the NAc of all rats after the behavioral paradigm. Therefore, we conducted another immunohistochemical staining using the Nef antibody and observed positive DAB staining in the area of infusion in both male and female Nef+cocaine rats compared to controls (Figure 4D, E). The Nef positive cells demonstrated a starlike morphology (shown with red arrows) around the anterior commissure, resembling astrocytes. To reduce the possibility that the sex-based behavioral outcomes could be due to the variability in Nef expression at the infused site, we analyzed the positive DAB staining and compared between males and females. No differences in Nef expression were observed between male and female tissues after the behavioral paradigm (Unpaired T-test=0.5454; Supplementary Figure S1).

For our behavioral analysis, we decided to separate male and female data because of differences in cocaine consumption between the sexes. Figure 5 shows male rat behavioral analysis during self-administration, extinction, cue-primed reinstatement, and cocaine-primed reinstatement, phases respectively. Data demonstrated that male rats expressing Nef had lower active (right) lever presses [days: 2 (p=0.0305), 6 (p=0.0005), 7 (p=0.0005), 8 (0.0483), 9 (p=0.0237), and 10 (p<0.0001)] and, consequently, fewer cocaine infusions [days: 7 (p=0.0067), 9 (p=0.0252), and 10 (p=0.0092)] during the self-administration sessions than cocaine male rats (Two-way ANOVA main treatment effect: Active Levers: F(3,31)=17.20, P=<0.0001; Infusions: F(3,31)=35.03, P=<0.0001; Session x treatment effect: Active Levers: F(69, 713)=6.892, P=<0.0001; Infusions: F(27,279)=6.067, P=<0.0001; Figure 5A, B). In addition, during cue-primed reinstatement, no differences were observed between cocaine-consuming groups (Two-way ANOVA main interaction: F(1,34)=0.003915, P=0.9505; Extinction vs. Reinstatement: F(1, 34)=67.35, P<0.0001; Cocaine vs. Nef+cocaine: F(1,34)=0.02510, P= 0.8751; Uncorrected Fisher’s LSD Post-Hoc: Reinstatement Cocaine vs Nef+cocaine: p=0.9464; Figure 5C). However, during cocaine-primed reinstatement, the Nef+cocaine group significantly increased cocaine seeking compared to the cocaine group (Two-way ANOVA main interaction: F(1,34)=0.1053, P=0.1053; Extinction vs. Reinstatement: F(1, 34)= 20.84, P<0.0001; Cocaine vs. Nef+cocaine: F(1,34)=3.472, P=0.0711; Uncorrected Fisher’s LSD Post-Hoc: Reinstatement Cocaine vs Nef+cocaine: p=0.0176; Figure 5D). This data suggests that Nef is causing higher cocaine-seeking behavior in male rats, and this may be due to higher sensitivity to cocaine when Nef is present in the NAc.

Similar results were obtained when exposing female rats to cocaine self-administration. However, during the self-administration phase, the cocaine and Nef+cocaine female groups had no differences in active lever presses or consumption of cocaine (Two-way ANOVA main treatment effect: Active Levers: F(3,33)=30.05, P=<0.0001; Infusions: F(3,32)=46.54, P=<0.0001; Session x treatment effect: Active Levers: F(69, 759)=5.871, P=<0.0001; Infusions: F(27,288)=4.618, P=<0.0001; Figure 6A, B). Female rats also demonstrated no difference in cocaine seeking during cue-primed reinstatement (Two-way ANOVA main interaction: F(1,36)=0.06235, P=0.8042; Extinction vs. Reinstatement: F(1, 36)=48.86, P<0.0001; Cocaine vs. Nef+cocaine: F(1,36)=0.1555, P= 0.6957; Uncorrected Fisher’s LSD Post-Hoc: Reinstatement Cocaine vs Nef+cocaine: p=0.6516; Figure 6C), but identical to male response, the female Nef+cocaine group increased the active lever presses compared to the female cocaine group during cocaine-primed reinstatement (Two-way ANOVA main interaction: F(1,36)=2.720, P=0.1078; Extinction vs. Reinstatement: F(1, 36)= 35.95, P<0.0001; Cocaine vs. Nef+cocaine: F(1,36)=4.246, P=0.0466; Uncorrected Fisher’s LSD Post-Hoc: Reinstatement Cocaine vs Nef+cocaine: p=0.0127; Figure 6D). This data once again suggests that Nef causes cocaine-seeking behavior when present in NAc astrocytes after a short access cocaine self-administration paradigm.

**Figure 6.**
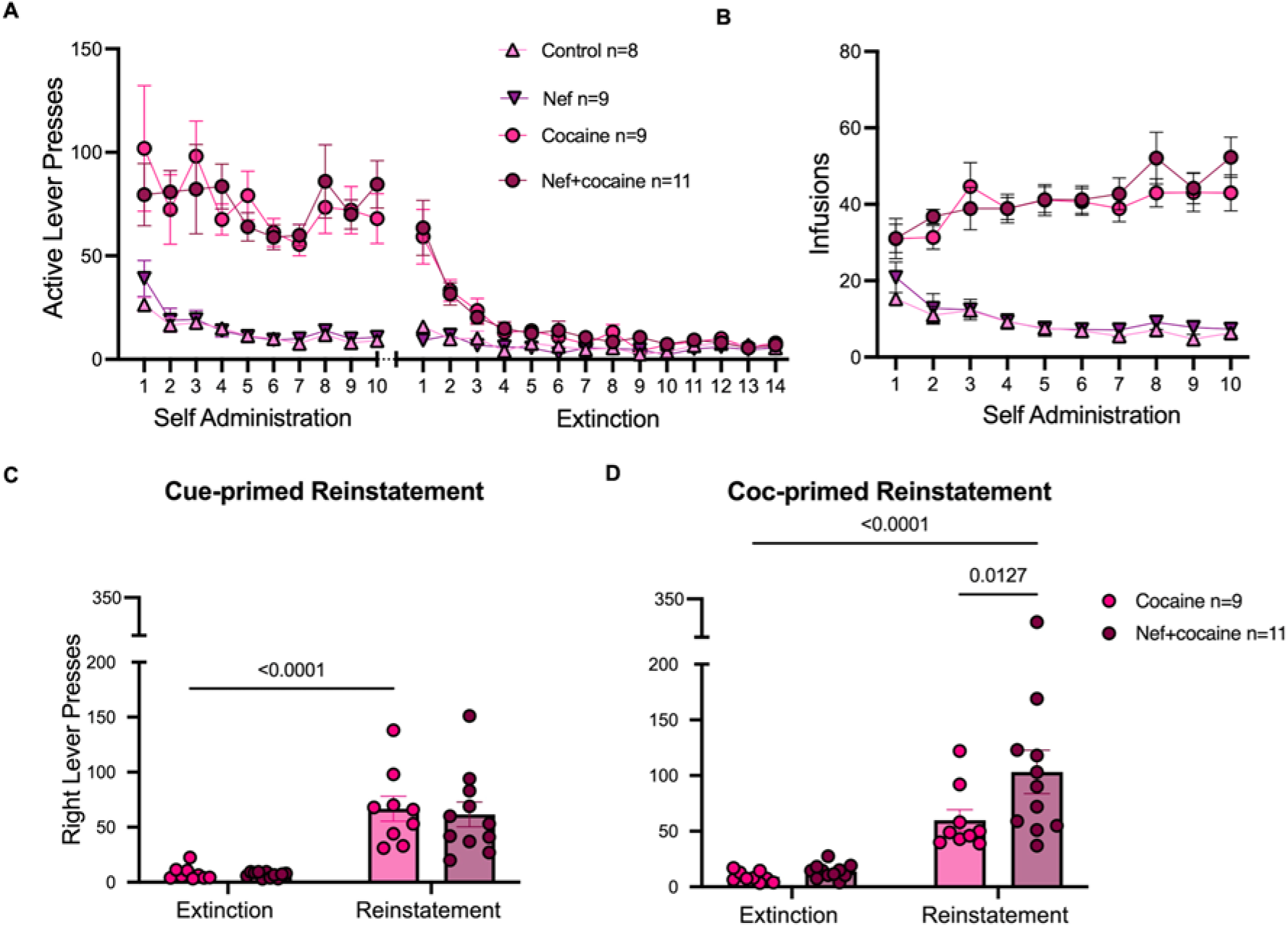
Nef-expressing female rats show more cocaine seeking behavior than Cocaine female rats. (**A**) Active lever presses of Control, Nef, Cocaine, and Nef+cocaine female rats during self-administration phase and extinction. (**B**) Cocaine infusions of Cocaine vs Nef+cocaine rats and saline infusions for Control and Nef rats during self-administration. No differences were observed between Cocaine and Nef+cocaine female groups. (**C**) Average right lever presses (correct) during extinction of Cocaine vs Nef+cocaine rats compared to right active lever presses during cue-primed reinstatement. (**D**) Average right lever presses of Cocaine vs Nef+cocaine animals compared to right active lever presses during cocaine-primed reinstatement. Nef+cocaine rats significantly increased active lever presses in cocaine primed reinstatement compared to Cocaine group. Two-way, repeated measures ANOVA indicated all rats reinstated cocaine seeking compared with extinction responding. Data represent mean +/- SEM.

### 3.3 Nef in the NAc causes sex-dependent differences in GLT-1 and GFAP after behavioral paradigm

After 24 hours of the cocaine-primed reinstatement, we sacrificed the rats for molecular analysis of glutamate transporters between each of the groups. Since the behavioral analysis was separated by sex, we decided to separate the protein analysis as well for this section. We collected NAc punches after sacrifice to quantify the protein expression of glutamate transporters GLT-1 and xCT. In this case, both in male and female rats, GLT-1 expression decreased with Nef and cocaine compared to the control group. However, we found differences in GLT-1 expression in the Nef+cocaine groups between males and females. In this case, male rats had similar GLT-1 expression with Nef, cocaine and Nef+cocaine, while females increased GLT-1 in the Nef+cocaine group, compared to Nef and cocaine alone [Males: One-way ANOVA main effect between treatments: F(3,31)=6.082, P=0.0022; Tukey’s Post-Hoc with multiple comparisons: Control vs Nef: p=0.0185; Control vs. Cocaine: p=0.0226; Control vs. Nef+cocaine: p=0.0017 (Figure 7A); Females: One-way ANOVA main effect between treatments: F(3,33)=8.325, P=0.0003; Tukey’s Post-Hoc with multiple comparisons: Control vs Nef: p=0.0159; Control vs. Cocaine: p=0.0023; Control vs. Nef+cocaine: p=0.9973; Figure 7D)]. On the other hand, both males and females decreased xCT expression with cocaine and Nef+cocaine, but no differences were observed with Nef alone [Males: One-way ANOVA main effect between treatments: F(3,31)=9.770, P=0.0001; Tukey’s Post-Hoc with multiple comparisons: Control vs Nef: p=0.1302; Control vs. Cocaine: p=0.0013; Control vs. Nef+cocaine: p=0.0001 (Figure 7B); Females: One-way ANOVA main effect between treatments: F(3,33)=7.114, P=0.0008; Tukey’s Post-Hoc with multiple comparisons: Control vs Nef: p=0.5485; Control vs. Cocaine: p=0.0009; Control vs. Nef+cocaine: p=0.0164 (Figure7E)], which demonstrated that xCT downregulation is driven by cocaine. We also analyzed GFAP expression after the behavioral paradigm to observe if astrogliosis was still present after 10 weeks with the presence of Nef as observed in our previous results. Interestingly, GFAP only trended towards an increase in the female Nef+cocaine group compared to control, but no differences were found between any of the other groups in male or female rats [Males: One-way ANOVA no main effect between treatments: F(3,31)=0.7416, P=0.5354; Tukey’s Post-Hoc with multiple comparisons: Control vs Nef: p=0.5866; Control vs. Cocaine: p=0.9960; Control vs. Nef+cocaine: p=0.7617 (Figure 7C); Females: One-way ANOVA no main effect between treatments: F(3,33)=2.385, P=0.0869; Tukey’s Post-Hoc with multiple comparisons: Control vs Nef: p=0.5599; Control vs. Cocaine: p=0.7915; Control vs. Nef+cocaine: p=0.0629 (Figure7F)]. This data suggests an involvement of GFAP expression in the modulation of glutamate transporters, specifically on an apparent neurotoxic environment. Overall, our data suggests that Nef disrupts GLT-1 transporter expression in the NAc and that its regulation is potentially involved in the observed cocaine-seeking behavior during the reinstatement phase.

**Figure 7.**
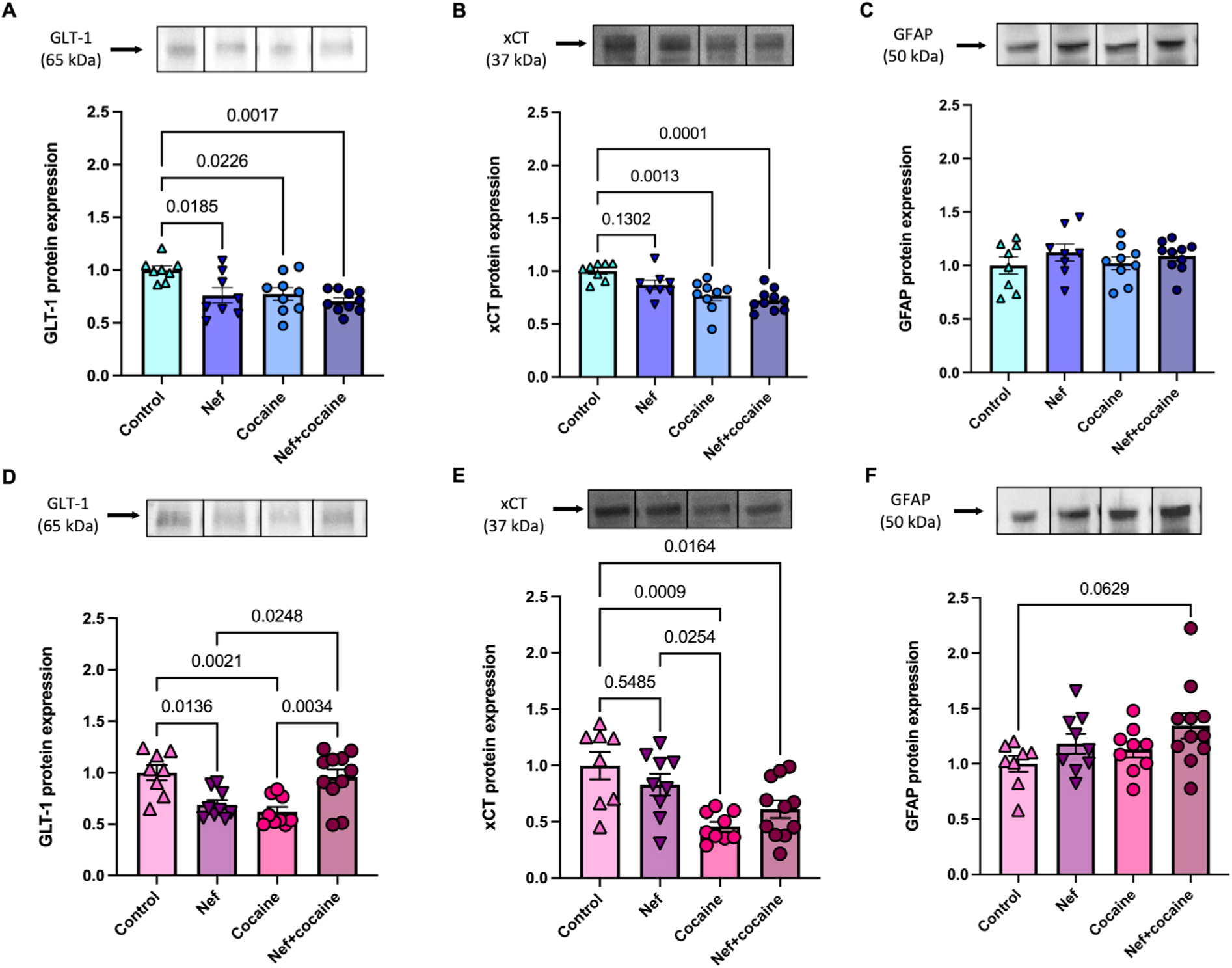
Nef alters glutamate transporter expression after cocaine self-administration paradigm in male and female rats. (**A**) Representative protein bands of GLT-1 (kDa) with analysis of western blots per group. Protein expression of GLT-1 shows decrease in all treatments compared to cocaine after cocaine-primed reinstatement of male rats. (**B**) Representative protein bands of xCT (37kDa) in NAc of male rats with analysis per group. Protein expression of xCT in the NAc shows decrease in cocaine and Nef+cocaine groups after behavioral paradigm of male rats. (**C**) Representative protein bands of GFAP (50kDa) in the NAc of male rats with analysis per group. GFAP expression shows no difference between groups of male rats after behavioral paradigm. (Male groups Control: n=8; Nef: n=8; Cocaine: n= 9; Nef+cocaine: n=10). (**D**) Representative protein blots of GLT-1 in the NAc of female rats with analysis per group. Protein expression of GLT-1 shows decrease in Nef and cocaine groups but increase in Nef+cocaine group after cocaine-primed reinstatement in female rats. (**E**) Representative blots of xCT in the NAc of female rats and analysis after behavioral paradigm per group. Protein expression of xCT in the NAc shows decrease in cocaine and Nef+cocaine after behavioral paradigm in female rats. (**F**) Representative images of GFAP protein expression with analysis per group of female rats. GFAP trends towards an increase in the Nef+cocaine female group compared to control 24 hours after cocaine-primed reinstatement. One-way ANOVA with Tukey Post-Hoc and multiple comparisons was used to compare data within groups. (Female groups Control: n=8; Nef: n=9; Cocaine: n=9; Nef+cocaine: n=11). Data represent mean +/- SEM adjusted to 𝛽𝛽-actin and normalized to control.

## 4. Discussion

Despite decades of research, our understanding of the underlying mechanisms driving drug-seeking in SUD remains limited. It is even less clear how the presence of HIV and viral proteins modifies these physiological alterations in the context of SUD in HIV infection. Although antiretroviral therapy (cART) manages to suppress HIV replication, astrocytes act as reservoirs in the brain with latent infection expressing HIV viral proteins, particularly Nef, resulting in inflammation and causing neuronal damage in the CNS [17, 37, 38]. Additionally, cocaine is known to accelerate the course of HIV infection, increasing viral neurotoxicity by astrocytes and ultimately disrupting neurotransmitter balance in the CNS [13, 39, 40]. While cocaine directly affects dopaminergic transmission, glutamatergic transmission also plays a significant role in mediating drug-seeking and relapse [41–43]. The glutamatergic system plays a significant role in neuroplasticity, drug sensitization, habit formation, and reinforcement learning [41], so our study focuses on glutamatergic effects. Producing HIV-1 neurotoxins in glial cells can alter glutamate homeostasis [40]. Despite this, there is a lack of research regarding the possible role of glutamate at the intersection of HIV-1 Nef and cocaine use disorder. Therefore, we aimed to address the relationship that exists between HIV-1 Nef, glutamate homeostasis, and cocaine use disorder in the NAc, a critical brain region associated with motivation, reward, and addiction.

In the present study, we assessed the effects of Nef after a single exposure of cocaine or a short-access, long-term exposure to cocaine (behavioral model) in the NAc of rats’ brains. Our results demonstrate several important findings. First, we show that expression of GLT-1 decreases in the NAc after one high-dose exposure to cocaine or astrocytic Nef expression in both male and female rats (Figure 2). The same was found by the end of the behavioral paradigm where rats self-administer cocaine repeatedly across 10 days (Figure 7). Interestingly, after the self-administration model, we found sex-based differences in GLT-1 transporter expression when exposed to Nef+cocaine. Male rats significantly decreased GLT-1, while the female group was unchanged compared to controls despite the significant reduction found for Nef or cocaine alone (Figure 7). Second, we demonstrate that multiple exposures to cocaine by short access self-administration increase cocaine-primed reinstatement in a Nef-treated Sprague Dawley rat model with no effect of sex (Figures 5 and 6). This data suggests that HIV-1 Nef or cocaine, modulates GLT-1 expression in the NAc, while their combination manifests differently in male or female rats (downregulated and unchanged, respectively). These molecular findings do not result in behavioral differences in reinstatement influenced by sex as we found that Nef increased cocaine-primed reinstatement in all rats but did not alter cue-primed reinstatement (Figures 5 and 6).; Interestingly, Nef expression moderated active lever pressing during acquisition only in male rats, hinting at an increased cocaine sensitivity (Figure 5). One limitation is that we could not assess GLT-1 expression during acquisition, preventing us from addressing the exact mechanism of GLT-1 and its role in the observed sex differences in lever pressing during this phase.

### 4.1 Astrocytic Nef and single dose of cocaine independently reduce GLT-1, but not xCT

The glutamate transporter EAAT2, or GLT-1 in rodents, is one of the most abundant proteins in the CNS and is pivotal in maintaining the balance between glutamate release and uptake [34]. It is responsible for 90% of extracellular glutamate reuptake by astrocytes, protecting against excitotoxicity [34, 44]. System Xc- is the primary source of extracellular glutamate in a variety of rodent brain areas [45], and it is responsible for up to 60% of extracellular glutamate in the NAc core [36]. The light chain, known as xCT, is primarily responsible for the function of the antiporter [46]. Cocaine use causes changes in extracellular glutamate and glutamate release in different structures associated with the reward system, specifically in the NAc. The NAc is one of the most critical structures to be involved in cocaine seeking. It is composed of glutamate terminals from neurons located in other reward-related areas, such as the thalamus (Th), amygdala (AMY), hippocampus (HIPP), and prefrontal cortex (PFC) [47]. Therefore, changes in glutamate levels in the NAc have been widely studied during different stages of cocaine administration, including self-administration and single/repeated cocaine doses with/without withdrawal. Reid and colleagues determined that a single high dose of cocaine (15-30mg/kg i.p) produces an increase in NAc extracellular glutamate of drug-naïve rodents. However, lower doses of cocaine (10mg/kg i.p. or less) do not acquire the same effect [48, 49]. In our study, we successfully demonstrated that one high dose of cocaine (15mg/kg i.p.) is sufficient to induce physiological responses in the NAc leading to downregulated protein expression of GLT-1 (Figure 2A, C, D); Nef only showed significant downregulation in immunofluorescent cells (Figure 2C). No significant changes were observed in xCT protein expression for any treatments (Figure 3). A possible explanation for the cells to moderate one transporter over the other under these conditions is that more than one cocaine dose is necessary to cause a significant alteration to all transporters and receptors associated with changes in glutamate levels. Also, Nef may not directly affect xCT in astrocytes. More studies should be done to test the mechanism through which xCT and GLT-1 levels are regulated by Nef and cocaine to better understand how changes may relate to behavior (as seen with cocaine-primed reinstatement in the behavioral model).

On the other hand, there are no studies done testing glutamate clearance or uptake within an HIV transgenic (Tg) or HIV-1 protein animal model with cocaine. It is known that HIV-1 is associated with neuroinflammation caused by multiple cytokines, such as IL-1B and TNF-a, which can contribute to a reduction in glutamate transporter expression [50, 51]. A study in multiple astrocytic cell lines demonstrated that Nef expression led to lower levels of glutamate uptake and release and a higher level of glutamate in the culture media of the infected cells [52]. However, they did not assess glutamate transport expression. For the first time, we demonstrated that astrocytic Nef expression in vivo caused a decrease in GLT-1 expression in the NAc (Figure 2A, C) compared to control rats. We also observed that one high dose of cocaine (15mg/kg i.p.) was enough to decrease the expression of this glutamate transporter after 24 hours. The downregulation observed could explain the excess extracellular glutamate detected in previous studies. However, our original hypothesis was that astrocytic Nef would exacerbate the rewarding effects of cocaine in the NAc. In this case, we did not observe any differences between Nef and cocaine treatments; when combined (Nef+cocaine), no change was observed when compared to the control group. A limitation of our study is that we could only assess GLT-1 at the time of sacrifice and so we do not know how the levels varied throughout the weeks of Nef expression, cocaine exposure and behavioral training. That was beyond the scope of this study and so it is challenging to speculate as to why we did not observe additive effects. It is possible that the already reduced GLT-1 expression caused by Nef or cocaine alone is already at its minimum capacity, and an exacerbated downregulation would result in neurotoxicity or cell death.

### 4.2 GLT-1 expression is reduced by Nef and cocaine independently after cocaine-primed reinstatement, while xCT downregulation is cocaine dependent

Next, we sought to discover what effects astrocytic Nef would cause in the context of cocaine-seeking behavior. Both male and female rats exhibited a higher cocaine-primed reinstatement with Nef in the NAc (Figures 5 and 6), findings which are consistent with those reported in HIV rodent models [53, 54]. Since the significant differences that we observed were during the cocaine-primed reinstatements, in our study, we decided to focus on the differences in glutamate transporter expression after re-exposure to cocaine. When astrocytic Nef was present in the NAc of the rats, GLT-1 transporter expression was also decreased compared to the control group after the behavioral paradigm (Figure 7). Several preclinical studies have revealed a reduction in GLT-1 expression in the NAc, more precisely in the NAc core, following brief (1 day) and prolonged (40–45 days) periods of cocaine abstinence, along with two to three weeks of extinction, from a typical (2 hours per day) cocaine self-administration [55–58]. Cocaine on its own can also reduce levels of xCT, especially in the NAc of rats after self-administration [35, 36]. According to some studies, downregulating these transporters supports drug-seeking behavior for cocaine and other addictive substances by causing glutamate spill-over in the NAc core [26]. Additionally, xCT expression is decreased in the NAc core and shell with relapse to cocaine-conditioned place preference (CPP) [59]. To date, no studies have been done to assess the effects of Nef and cocaine on glutamate transporter expression. Our study observed differences in xCT expression between cocaine and Nef-treated male and female rats, where the only effect that occurred was cocaine driven after the behavioral paradigm. Our results also demonstrate that Nef can reduce GLT-1 expression in astrocytes and that this reduction is long-lasting in the NAc. We believe that one possible mechanism that could modulate glutamate homeostasis in the presence of Nef is by reducing β-catenin, a regulator of glutamate transporters in astrocytes [60, 61]. Nef’s structure aligns with the β-catenin central binding motif, allowing it to interact with β-catenin directly through a comparable region at its C-terminus [61]. It is unknown, nevertheless, the effect HIV-1 Nef protein has on β-catenin-regulated genes, like GLT-1. Other studies have also mentioned that the production of Nef by astrocytes triggers the release of tumor necrosis factor alpha (TNF-α) from monocyte-derived macrophages (MDMs) and CD4-T cells, as well as an increase in the pro-inflammatory mediator interleukin (IL)-1β [38, 62, 63]. TNF-α and IL-1β can decrease glutamate transporter expression, which leads to an imbalance in glutamate homeostasis and neuroinflammation caused by HIV-1 [50, 51]. These are possible ways Nef could be involved in the decreased expression of glutamate transporters after the self-administration paradigm. Cocaine alone has also been seen to result in β-catenin induced changes after abstinence of chronic doses, where 3 hours and 24 hours after the last injection caused an increase in β-catenin in the NAc core of male rats that specifically showed behavioral sensitization [64]. This data suggests that Nef and cocaine may have an interconnected effect in the Wnt/ β-catenin pathway, which could result in a potential therapeutic target for future treatments in this population. However, more studies should be done to assess the mechanism by which HIV-1 Nef may be involved in the downregulation of glutamate transporters in the context of HIV and cocaine use disorder.

### 4.3 Nef and cocaine in combination show sex dependent effects on GLT-1 expression after behavioral paradigm

Moreover, we observed a sex difference in our molecular data after the behavioral paradigm. Male rats showed a decrease in GLT-1 expression with Nef+cocaine 24 hours after cocaine primed reinstatement, while the female rats exhibited an increase compared to the cocaine alone group. This difference may be influenced by estrogen receptors expressed on astrocytes. Estrogens can bind to and activate estrogen receptors (ERs) on astrocytes [65] and can impact astrocyte activity in the brain. Specifically, estradiol, a potent estrogen hormone, has been linked to affecting glutamate metabolism in the brain and plays a protective role against neurodegeneration [66–68]. Although done in cultured astrocytes, researchers have found that estrogen levels positively increase glutamate transporter expression of GLT-1 and GLAST on mRNA and protein levels, decreasing extracellular glutamate levels [66, 69]. These results may help explain the increase in GLT-1 expression observed in the NAc of our female rats after the behavioral paradigm, as a result of a compensatory mechanism to protect against neurotoxicity. However, the decrease in GLT-1 specifically observed in the Nef+cocaine male rats is thought to be due to sensitivity to the drug. Even if these results were expected, we did not anticipate a behavioral difference during the self-administration phase. Interestingly, even if Nef+cocaine male rats consumed less than the cocaine group during the acquisition phase, both acquired a similar cue-induced reinstatement after an extinction period. Ample evidence suggests that GLT-1 is downregulated after different withdrawal periods and that a reduction in this transporter, specifically in the NAc, is associated to stronger neuroadaptations linked to cocaine seeking or relapse [55, 70, 71]. Our Nef+cocaine male animals not only consumed less cocaine than the cocaine group, but they later demonstrate a stronger cocaine induced seeking behavior. This data suggests that Nef may be inducing higher sensitivity to cocaine that is only observed in male rats.

## 5. Conclusions

Our study provides insights into the complex interactions between HIV-1 Nef, cocaine, and glutamate homeostasis in the NAc of rats. We found that both astrocytic Nef and cocaine exposure independently decreased the expression of GLT-1, with notable sex-based differences observed after multiple doses. Male rats exhibited a significant reduction in GLT-1 expression when exposed to Nef and cocaine, while females showed an increase, suggesting an estrogen-mediated neuroprotection. Behaviorally, Nef in the NAc heightened cocaine-primed reinstatement in both sexes, indicating a heightened vulnerability to relapse. These findings highlight the essential role of HIV-1 Nef and cocaine in glutamate dysregulation, which can relate to higher cocaine use by patients with HIV. Some limitations in our methodology were brought up during the study. First, we detected a low signal for Nef. That may be due to low infection or sensitivity and specificity of the Nef antibody (a known problem in the field). The percentage of Nef expression can be close to 0.1% in astrocytes positive for HIV [72]. A low expression of the protein is expected, which more closely mimics how the actual virus interacts with humans [73]. Our study still detected differences between treatments with low Nef expression. However, we still do not know how Nef’s downregulation of glutamate transporter expression in astrocytes involves neuronal changes in the NAc. We understand that focusing on one neurotoxin instead of the whole virus is also a limitation. Clinically, cART effectively lowers HIV viral load to the extent of non-detection, yet no antiretrovirals specifically block HIV transcription. Our selection of Nef, an early HIV-1 protein and a potent neurotoxin, builds on our prior work demonstrating that limited HIV-1 expression is sufficient to cause learning impairment [33] and now is shown to alter cocaine seeking behavior. We note that CNS viral reservoirs persist, raising the prevalence of HAND in individuals with long-term infection. In patients suffering from severe HAND, up to 19% of astrocytes test positive for HIV-1. The degree of neuropathological alterations is correlated with the level of infection [18]. Thus, it is essential to determine whether these reservoirs, even when restricted from replicating, may still express viral neurotoxins like Nef that are crucial to the mechanisms behind this comorbidity. We also did not focus on how changes in the NAc could alter or be altered by neuronal projections in other areas of the reward system, such as the PFC, HIPP, Th, and AMY. Future studies should focus on understanding the mechanism through which HIV-1 Nef disrupts glutamate homeostasis, ultimately altering synaptic plasticity in the context of HIV and cocaine substance use disorder.

## Supplementary Materials

**Figure S1:**
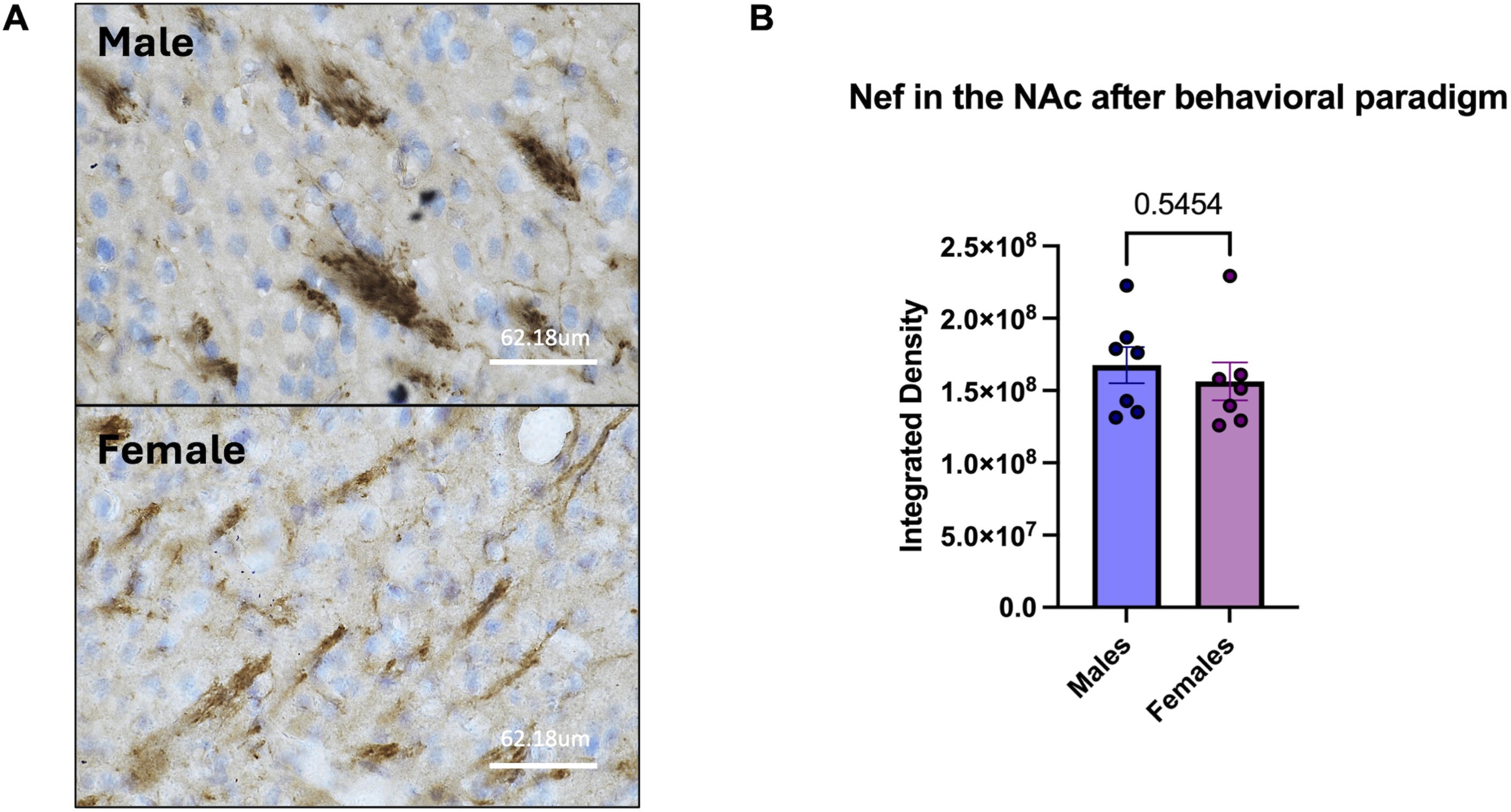
Quantified Nef expression in NAc between male and female tissues after behavioral paradigm. (A) Representative images of immunohistochemical staining with Nef antibody (1:50) in NAc slice of male (top) and female (bottom) rat Nef treated brain tissue. DAB staining in brown with hematoxylin for nuclear counterstain. Pictures of 30um thick tissues taken at 60x magnification (62.18um scale). (B) Integrated density of Nef expression in the NAc shows no significant differences between males and females. Each sample represents an average of 4-5 60x mag fields per rat in each treatment. Unpaired T-test was used to compare data within groups (Males: n==7; Females: n==7).

## Author Contributions

Conceptualization, J.P.-T., R.J.N.; methodology, J.P.-T., B.V.-P., M.C.-R., M.S.-O., and R.J.N.; validation, J.P.-T., M.C.-R., M.S.-O., and R.J.N.; formal analysis, J.P.-T., B.V.-P., M.S.-O. and R.J.N.; investigation, J.P.-T., B.V.-P., Y.M.-B., M.C.-R.; resources, M.S.-O., and R.J.N.; data curation, J.P.-T.; writing—original draft preparation, J.P.-T., Y.M.-B. and R.J.N.; writing—review and editing, J.P.-T., Y.M.-B., M.S.-O., and R.J.N.; funding acquisition, M.S.-O. and R.J.N. All authors have read and agreed to the published version of the manuscript.

## Institutional Review Board Statement

The animal study protocols were approved by the Animal Care and Use Committee (IACUC) of Ponce Health Sciences University Institutional (protocol codes: 2006000298, approved 27 July 2020; 2307143428, approved 07 August 2023).

## Informed Consent Statement

Not applicable.

## Data Availability Statement

The raw data supporting the conclusions of this article will be made available by the authors on request.

## Acknowledgments

A special thanks to Maria Colon-Romero, Jaydie Valles-Ortiz, and Ursula Gelpi-Dominguez, for their assistance at different stages of this project. We also acknowledge support from PHSU/PRI Biomedical Sciences Program Facilities including the Molecular and Genomics Core, BRAIN Core and Protein Core.

## Disclaimer/Publisher’s Note

The statements, opinions and data contained in all publications are solely those of the individual author(s) and contributor(s) and not of MDPI and/or the editor(s). MDPI and/or the editor(s) disclaim responsibility for any injury to people or property resulting from any ideas, methods, instructions or products referred to in the content.

